# Adaptation to binocular anticorrelation results in increased neural excitability

**DOI:** 10.1101/549949

**Authors:** Reuben Rideaux, Elizabeth Michael, Andrew E Welchman

## Abstract

Throughout the brain, information from individual sources converges onto higher order neurons. For example, information from the two eyes first converges in binocular neurons in area V1. Many neurons appear tuned to similarities between sources of information, which makes intuitive sense in a system striving to match multiple sensory signals to a single external cause, i.e., establish causal inference. However, there are also neurons that are tuned to dissimilar information. In particular, many binocular neurons respond maximally to a dark feature in one eye and a light feature in the other. Despite compelling neurophysiological and behavioural evidence supporting the existence of these neurons (Cumming & Parker, 1997; Janssen, Vogels, Liu, & Orban, 2003; Katyal, Vergeer, He, He, & Engel, 2018; Kingdom, Jennings, & Georgeson, 2018; Tsao, Conway, & Livingstone, 2003), their function has remained opaque. To determine how neural mechanisms tuned to dissimilarities support perception, here we use electroencephalography to measure human observers’ steady-state visually evoked potentials (SSVEPs) in response to change in depth after prolonged viewing of anticorrelated and correlated random-dot stereograms (RDS). We find that adaptation to anticorrelated RDS results in larger SSVEPs, while adaptation to correlated RDS has no effect. These results are consistent with recent theoretical work suggesting ‘what not’ neurons play a suppressive role in supporting stereopsis (Goncalves & Welchman, 2017); that is, selective adaptation of neurons tuned to binocular mismatches reduces suppression resulting in increased neural excitability.

## INTRODUCTION

It remains an important challenge in neuroscience to understand how the brain combines a pair of two-dimensional retinal images to support 3D perception. Classically, this problem has been framed as one of matching features between the two eyes, i.e., solving the “stereoscopic correspondence problem”, so that depth of objects can be triangulated (Julesz & Chang, 1976; Marr & Poggio, 1976). This problem is nontrivial, as the number “false matches” (i.e., correspondences between features that do not originate from the same object) rapidly increases with the number of to-be-matched elements.

Random-dot stereograms (RDS) are frequently used to investigate binocular vision because of their ability to divorce information about 2D form from differences between the two eyes. These stimuli are composed of many self-similar features, potentially posing a severe challenge to establishing binocular correspondence. The classic framework for understanding stereopsis is to find correspondence by considering a range of potential disparities and selecting the offset that maximises the image similarity between the two eyes (Fleet, Wagner, & Heeger, 1996; Ohzawa, DeAngelis, & Freeman, 1990). This makes intuitive sense, however, many disparity-selective neurons in V1 appear poorly optimised for such a computation in that they respond maximally to different images presented on the two retinae (Cumming & Parker, 1997; Read & Cumming, 2007). Moreover, binocular neurons can show tuning to images that are difficult to imagine being produced in the real world. A prime example of this is tests of neural function with anticorrelated RDSs (aRDS) in which the polarity of image features is reversed between the two eyes. Unlike correlated stereograms (cRDS), viewing aRDS does not support reliable depth perception; nevertheless, many disparity-selective neurons in V1 respond strongly to these stimuli. Despite empirical evidence supporting the existence of these neurons in macaques (Cumming & Parker, 1997; Janssen et al., 2003; Tsao et al., 2003) and humans (Katyal et al., 2018; Kingdom et al., 2018), their functional role remains opaque.

Recent theoretical work suggested a potential explanation for neurons tuned to mismatched binocular features. In their binocular likelihood model of stereopsis, Goncalves and Welchman (2017) suggested a simple decoding rule for binocular neurons: information about depth can be read out from a population of binocular neurons where the decoding scheme is based on the cross-correlation between the encoding receptive fields. Under this scheme, the activity of a binocular neuron can lead to increased excitation for a particular depth interpretation, or drive suppression of a specific depth estimate. By reading out a population of binocular neurons, it is possible to derive a likelihood estimate of the depth of the scene. This provides a plausible explanation for why neurons should respond to binocular correspondences that do not relate to a single physical object in the environment. In particular the ‘what not’ responses of binocular neurons can be used to drive suppression of unlikely interpretations of the scene. Despite this theoretical promise, there is little empirical evidence for the role of ‘what not’ responses in the human visual system.

The idea that binocular mismatches are used to drive suppression in visual cortex yields a distinct prediction concerning the balance of excitation and inhibition following a period of adaption. In particular, adapting the responses of units that drive suppression should lead to less inhibition, thereby increasing the net excitation of the cortex. To investigate the role of ‘what not’ responses within the visual cortex, here we use electroencephalography to measure human observers’ brain activity during and after prolonged viewing of aRDS. Specifically, we measure steady-state visually evoked potentials (SSVEP) in response to cRDS and aRDS, following adaptation to either aRDS or cRDS. We find that following adaptation to aRDS, SSVEP amplitude in response to cRDS increases relative to a pre-adaptation baseline. These results are consistent with the idea that ‘what not’ responses play a suppressive role in supporting stereopsis; that is, selective adaptation of ‘what not’ responses reduce suppression, resulting in increased neural excitability.

## METHODS

### Participants

Observers were recruited from the University of Cambridge, had normal or corrected-to-normal vision, and were screened for stereo deficits. 33 right-handed human adults (8 male; 25.2±4.8 years) participated in the main experiment, however, two were not included in the analysis: one was unable to see depth in the stimulus, and a hardware fault stopped acquisition mid-way through the experiment for the other. Of the 31 participants included in the analysis, 22 completed all experimental conditions while the remaining 9 did not participate in the baseline condition. 22 right-handed human adults (5 male; 25.4±4.6 years) participated in the control experiment. Experiments were approved by the University of Cambridge ethics committee; all observers provided written informed consent.

### Apparatus and stimuli

Stimuli were generated in MATLAB (The MathWorks, Inc., Matick, MA) using Psychophysics Toolbox extensions (Brainard, 1997; Pelli, 1997). Binocular presentation was achieved using a pair of Samsung 2233RZ LCD monitors (120Hz, 1680×1050) viewed through mirrors in a Wheatstone stereoscope configuration. The viewing distance was 50 cm and the participants’ head position were stabilized using an eye mask, headrest and chin rest. Eye movement was recorded binocularly at 1 kHz using an EyeLink 1000 (SR Research Ltd., Ontario, Canada).

Adaptation stimuli consisted of random dot stereograms (RDS; 12°×12°) on a mid-grey background surrounded by a static grid of black and white squares intended to facilitate stable vergence. Dots in the stereogram followed a black or white Gaussian luminance profile, subtending 0.07° at half maximum. There were 108 dots/deg2, resulting in ∼38% coverage of the background. In the centre of the stereogram, 4 wedges were equally distributed around a circular aperture (1.2°), each subtending 10° in the radial direction and 70° in polar angle, with a 20° gap between wedges (**Fig. 1a**). Dots comprising the wedges were offset by 10 arcmin between the left and right eyes, while the remaining dots had zero offset. Stimuli were presented for 1 s and separated by 1 s inter-stimulus-intervals consisting of only the background and fixation cross. On each presentation, we applied a random polar rotation to the set of wedges such that the disparity edges of the stimuli were in different locations for each stimulus presentation (i.e., a rigid body rotation of the four depth wedges together around the fixation point). Every eight presentations we reversed the sign of the disparity of the wedges (crossed and uncrossed; **Fig. 1c**). At a given timepoint, all wedges were presented the same disparity. In the centre of the wedge field, we presented a fixation square (side length = 1°) paired with horizontal and vertical nonius lines.

**Figure 1.**
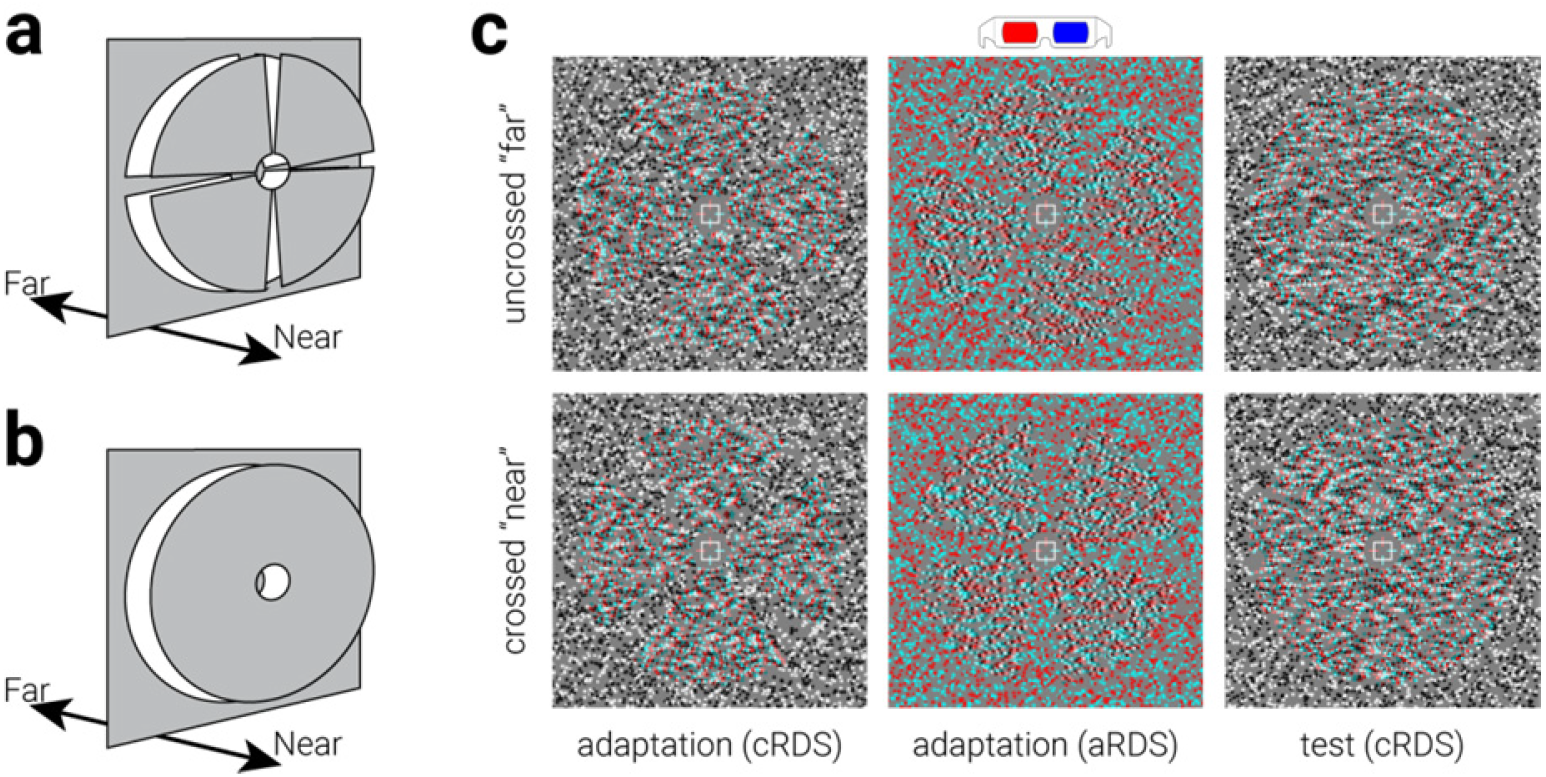
Adaptation and test stimuli used in the experiment. Diagram of the depth arrangement of the (**a**) adaptation and (**b**) test stimuli. (**c**) Example stimuli used in the experiment.

Test stimuli were similar to adaptation stimuli, except that instead of rotating wedges, an annulus was used (**Fig. 1b, c**). The depth sign of the annulus was reversed at 4 Hz frequency while all dots were regenerated at 20 Hz frequency.

### Procedure

Participants underwent three different runs: an initial baseline run, followed by correlated and anticorrelated runs (counterbalanced across participants). The baseline run consisted of 5 blocks of test stimulus presentations, each lasting 25 s, separated by 20 s blank inter-block-intervals. Correlated runs consisted of 12 adaptation blocks, each followed by a 2 s blank inter-block-interval prior to a test block. Adaptation blocks consisted of 64 s of adaptation stimuli presentations (32 presentations total), and test blocks were identical to those in the baseline run. Anticorrelated runs were identical to correlated runs, except that the polarity of all dots in the left eye was reversed during the adaptation blocks. During adaptation blocks, we instructed participants to fixate on the central fixation square while performing a Vernier discrimination task (Preston, Li, Kourtzi, & Welchman, 2008). During test blocks, we instructed observers to maintain fixation and limit blinks. Correlated RDS (cRDS) were used as the test stimulus for all conditions, providing an equal test of the effects of adaptation. While viewing either anticorrelated or correlated RDS can produce an electrophysiological response, cRDS were used as the test stimulus as they evoke a larger response than anticorrelated RDS (Braddick et al., 1980; Petrig, Julesz, Kropfl, Baumgartner, & Anliker, 1981), providing a better signal-to-noise ratio and thus a more sensitive measure of neural change resulting from adaptation.

### EEG

Electroencephalography data were acquired from all 33 participants with a 64 channel cap (BrainCap, Brain Products GmbH). Data were recorded using BrainVision Recorder software. Caps were fitted with 61 Ag/AgCl electrodes positioned according to the standard 10-20 system (Fp1 Fp2 F3 F4 C3 C4 P3 P4 O1 O2 F7 F8 T7 T8 P7 P8 Fz Cz Pz IO FC1 FC2 CP1 CP2 FC5 FC6 CP5 CP6 FT9 FT10 F1 F2 C1 C2 P1 P2 AF3 AF4 FC3 FC4 CP3 CP4 PO3 PO4 F5 F6 C5 C6 P5 P6 AF7 AF8 FT7 FT8 TP7 TP8 PO7 PO8 Fpz CPz POz Oz). Data were acquired with a reference electrode at FCz. Electro-oculograms were also acquired, using two pairs of bipolar electrodes placed horizontally and vertically around the left eye. Data were highpass filtered online at 0.1 Hz and acquired with a 1 Hkz sampling rate. Temporal markers were sent from the stimulus presentation computer to mark the onset of the stimulus. These timings were validated using a pair of photodiodes attached to the two stimulus presentation screens.

Preprocessing and analyses were performed in MATLAB, using the EEGLAB toolbox (Delorme & Makeig, 2004) and custom in-house scripts. Data were first filtered offline with a 1 Hz highpass and a 40 Hz lowpass filter. For the SSVEP analysis, each epoch was extracted around the test duration to include a period of 29 s (2 s prior to the first test-stimulus onset to 2 s after the offset of the final test-stimulus); thus, there were 12 epochs per adaptation condition and 5 in the baseline, all of which were included in the analysis. For the ERP analysis, epochs were extracted around the stimulus onset to include a 1 s pre- and 1 s post-stimulus period (384 epochs per adaptation condition). The post-stimulus period, therefore, did not include any data from the next stimulus presentation. Epochs used in the ERP analysis were visually inspected, and artefactual epochs were rejected (excluding eye movements). All data were re-referenced to an average reference across all channels, and then ICA decomposition was applied for the purpose of artefact identification. Resultant ICA components were visually inspected before rejection. Only components reflecting eye movements and other likely muscle artefacts were removed. These components were identified by characteristic features in the component time course and power spectrum, in addition to their frontal topography.

To equate the sensitivity of the baseline and adaptation SSVEP measurements, we matched the length of data in each calculation by including data from the first half of the adaptation epochs (4-13.5s), and 22.8s of data from the baseline epochs (2-24.8s). We avoided including data immediately from stimulus onset to avoid contamination by any onset evoked potential.

### Eye tracking

Owing to the bespoke experimental setup (i.e., recording eye position from behind one-way mirrors in a haploscope), the eye tracker would occasionally fail to track participants’ eyes for an entire block. In order to draw within-subjects comparisons, we only included the subjects for whom data was available for both experimental conditions in the eye tracking analysis (*n*=21). Prior to analysis, eye movement data was screened to remove noisy and/or spurious recordings. To remove spurious significant differences in the time course between conditions, a cluster correction was applied. Clusters were defined by the sum of their constituent (absolute) t-values and compared to a null hypothesis distribution of clusters produced by shuffling the condition labels (1000 permutations). Clusters below the 95^th^ percentile of the null hypothesis distribution were disregarded.

## RESULTS

### SSVEP analysis of test stimuli

If mismatches between the two eyes evoke inhibitory activity, we would expect that selectively reducing the responsiveness of the neural mechanisms that respond to mismatches (through adaptation) would lead to less inhibitory activity and thus relatively more excitability. To test this idea, observers were initially adapted to binocular mismatches by viewing anticorrelated random-dot stereograms (aRDS) for a prolonged period (64 s). Then, following adaptation, participants viewed a correlated RDS (cRDS) comprised of black and white blobs depicting an annulus that was either near or far relative to the background. We used the event-related potential (ERP) evoked by changing the depth sign of the annulus (from near to far or far to near) as an index of stereoscopic related activity and the ERP evoked by changing the luminance polarity of blobs (from white to black or black to white) as an index of non-stereoscopic related activity. Similar to previous work (Cottereau, McKee, Ales, & Norcia, 2011, 2012; Cottereau, McKee, & Norcia, 2012), we rapidly changed the stimuli at two frequencies, i.e., 4 Hz for depth sign and 20 Hz for luminance polarity, producing two distinct steady-state visually evoked potentials (SSVEPs). We measured the activity evoked by the stimulus changes by performing a Fourier transform on the data, converting it from the time domain to the frequency domain, and taking the signal-to-noise ratio (SNR) between the peak at the target frequencies (4 Hz & 20 Hz) and the baseline noise in the spectrum (from bins either side of the target frequency). For comparison, we also measured observers’ SSVEP in response to the test stimulus following adaptation to cRDS and without prior adaptation (baseline).

To guide electrode selection, we computed the 4 Hz SNR of each sensor during the baseline test run. As anticipated, we found the highest SNR for occipital and parietal sensors (**Fig 2a**). Based on these results, and previous electrophysiological work on binocular disparity (Cottereau et al., 2011; Cottereau, McKee, & Norcia, 2012), we included all parietal and occipital (i.e., Oz, O1, O2, Poz, PO3, PO4, PO7, PO8, Pz, P1, P2, P3, P4, P5, P6, P7, & P8) sensors in the main SSVEP analysis.

**Figure 2.**
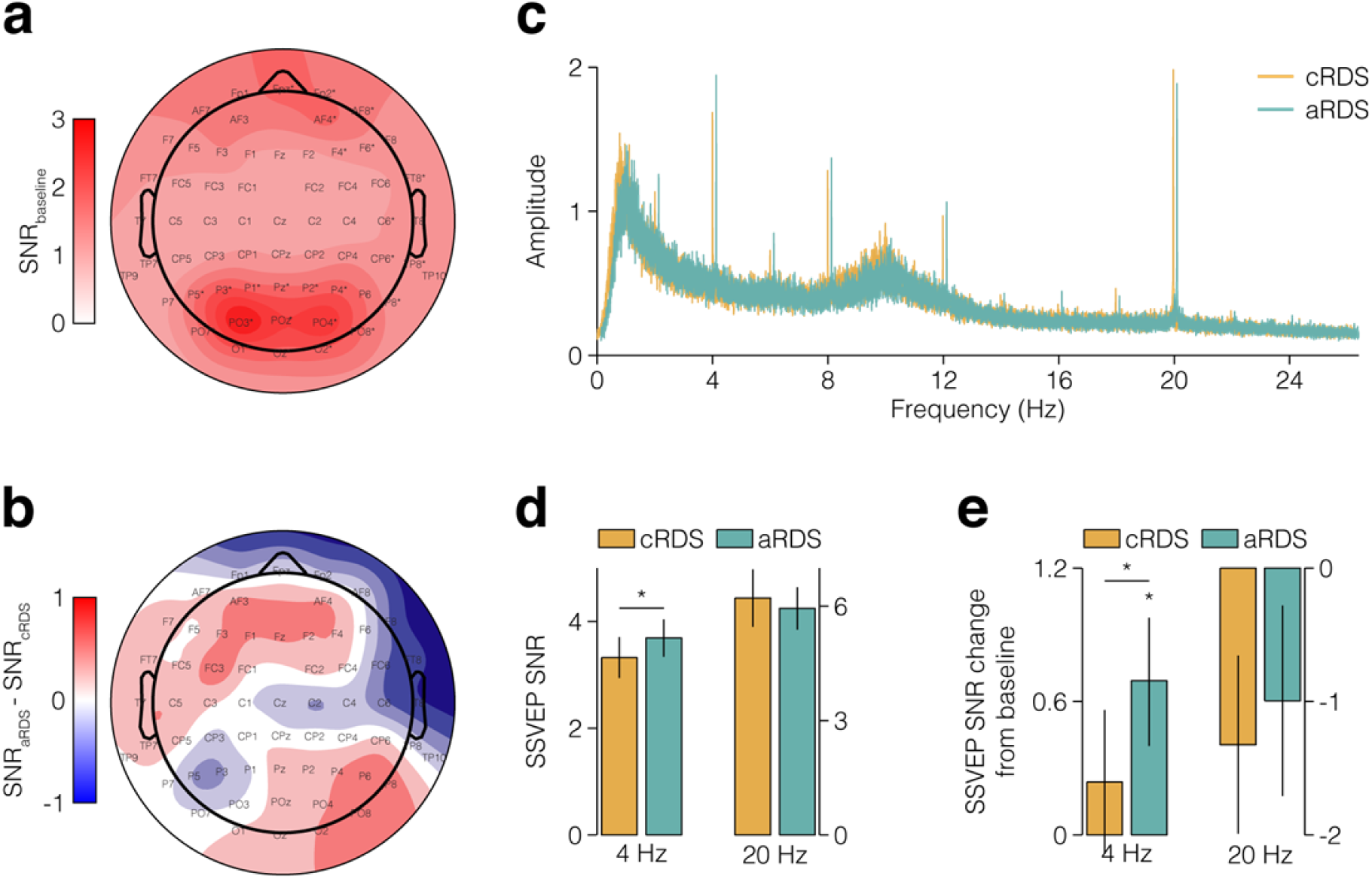
Steady-state visually evoked potential (SSVEP) in response to the test stimulus. (**a**) Topographic map showing the SSVEP signal-to-noise ratio (SNR) in response to the test stimulus without prior adaptation. (**b**) Same as (**a**) but for the difference in response to the test stimulus following adaptation to either correlated or anticorrelated random-dot stereograms (RDS) across. (**c**) The SSVEP SNR response spectra (averaged across parietal and occipital sensors) to the test stimulus following adaptation to c/aRDS, for all participants (*n*=31). The aRDS spectrum is horizontally offset to facilitate comparison with the cRDS spectra. (**d**) Same as (**c**) but isolating the depth alternation (4 Hz) and dot refresh (20 Hz) frequencies. A repeated-measures analysis of variance (ANOVA) of SSVEP SNR revealed a main effect of frequency (4/20 Hz; *F*_1,30_=10.47, *P*=.003) and an interaction between frequency and adaptation (c/aRDS; *F*_1,30_=4.23, *P*=.049), but no main effect of adaptation (*F*_1,30_=0.01, *P*=.94). For subjects who completed the baseline measurement (*n*=15), (**e**) shows the same as (**d**) but referenced to baseline SSVEP amplitude. Asterisks in (a) indicate sensors with SNR significantly than >1 following Bonferroni correction (N=64, α = 7.8e^−8^); asterisks in (d, e) indicate significant differences. Error bars indicate s.e.m.

Overall, we observed differences in activity in response to the test following adaptation to cRDS and aRDS **(Fig 2b, c).** To test the prediction that prolonged viewing of binocular mismatches increases neural excitability we pooled SSVEP SNR across parietal and occipital sensors and compared responses at the depth-change frequency (4 Hz). In line with the prediction, we found that SSVEP SNR was significantly higher following adaptation to aRDS compared to cRDS (all subjects: *t*_30_=2.49, P=.019; subjects with baseline: *t*_21_=2.77, *P*=.011; Fig 2d). A possible concern is that the difference in SSVEP SNR between adaptation conditions was due to a decrease in excitability following adaptation to cRDS, rather than an increase following adaptation to aRDS. However, we found no evidence for this: while adaptation to aRDS significantly increased SSVEP SNR relative to baseline (t_21_=2.39, P=.026), adaptation to cRDS produced no significant change (t_21_=0.74, P=.469; **Fig 2e**); these results are consistent with the interpretation that adaptation to binocular mismatches increased neural excitability.

Another possible concern is that the increased excitability we observed in the primary visual cortex following adaptation to aRDS is not specific to stereopsis, but a generalized effect. However, we found no evidence for a difference in SSVEP amplitude at the frequency of the dot refresh (20 Hz) between aRDS and cRDS adaptation conditions (paired t-test, t_30_=1.07, P=.294; **Fig 2d, e**), suggesting that the effect relates to changes in depth and not luminance. Another possible concern is that the effect might be driven by a subset of participants with low SSVEP amplitude across all conditions. Specifically, SSVEP amplitude measurements are less reliable at low values, thus a subset of unreliable values could potentially yield a false positive. However, while we found significant correlations between baseline, cRDS, and aRDS SSVP amplitude (Pearson correlation, subjects with baseline: *n*=22, baseline-cRDS: *r*=.489, P=.021, baseline-aRDS: *r*=.596, *P*=.003, cRDS-aRDS: r=.897, P=1.5e^−13^; all subjects: cRDS-aRDS: *r*=.92, P=1.5e^−13^; **Fig 3a-c**), we found no evidence of a relationship between baseline amplitude and the difference between aRDS and cRDS (Pearson correlation, *n*=22, *r*=.219, *P*=.327; **Fig 3d**).

**Figure 3.**
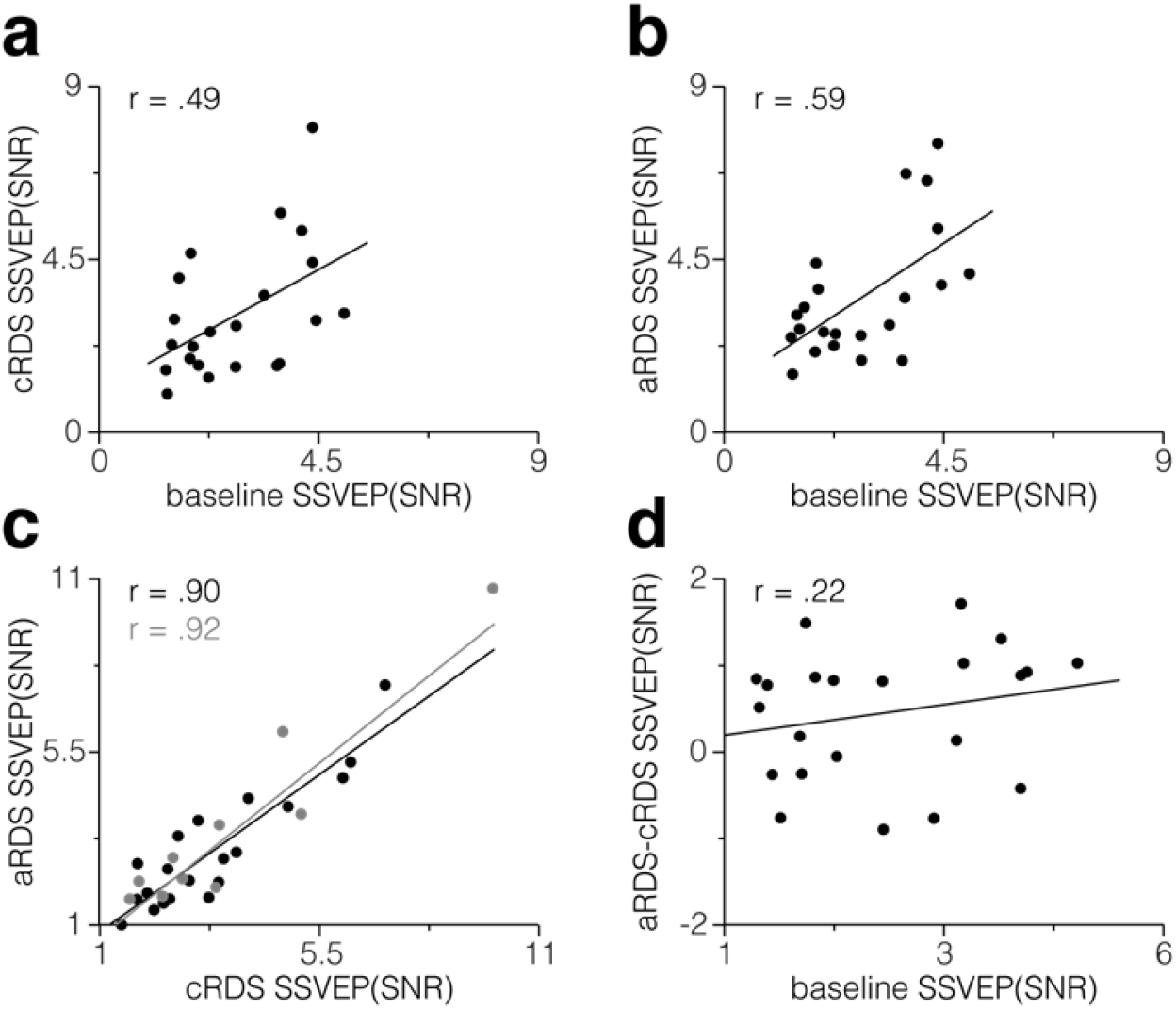
Relationship between steady-state visually evoked potential (SSVEP) signal-to-noise (SNR) across conditions. (**a**) SSVEP SNR measured in the response to the test stimulus following cRDS adaptation as a function of SSVEP SNR without adaptation. (**b**) Same as (**a**) but following aRDS adaptation. (**c**) Same as (**b**) but as a function of SSVEP SNR following cRDS adaptation. Black and grey dots indicate participants who did and didn’t complete the baseline condition, respectively. Black and grey lines indicate the least squares regression not including (*n*=15) and including (*n*=24) participants who completed the baseline, respectively. (d) The difference in the effect of cRDS and aRDS adaptation on SSVEP SNR as a function of SSVEP SNR without adaptation.

Finally, another possible explanation for the effect might be that observers’ attentional allocation during the adaptation period varied between conditions. However, we found no evidence for a difference in performance on the attentionally demanding Vernier task between conditions (paired t-test, *t*_30_=1.22, P=.23). Similarly, we found no evidence for a difference in eye movements in response to either the adaptation or test stimuli (**Fig 4a, b**).

**Figure 4.**
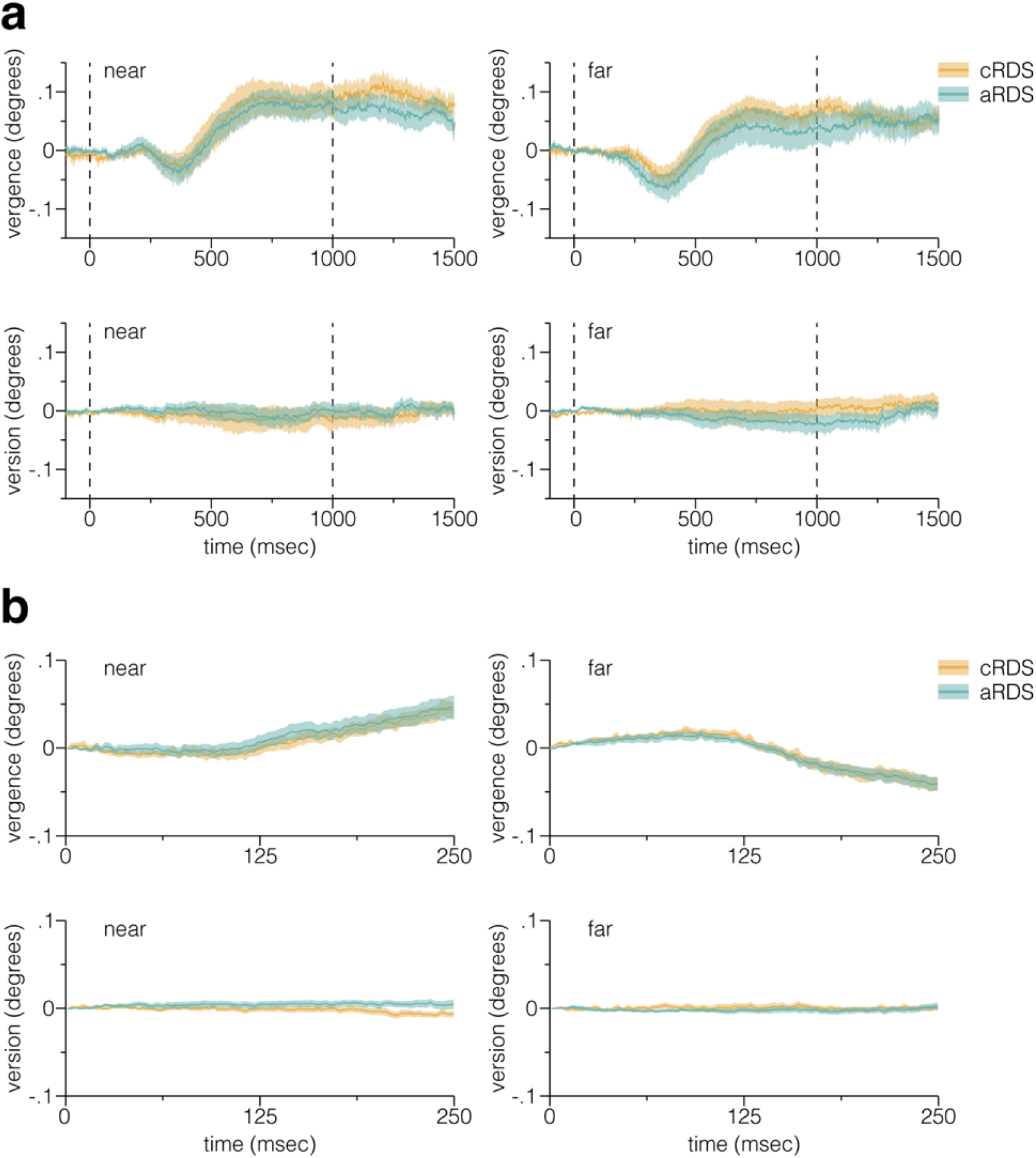
Comparison of eye movements between (c/aRDS) adaptation conditions. We assessed whether the adaptation effects observed could be explained by differences in eye movements during stimulus presentation, by comparing observers’ eye position during the adaptation and test periods between cRDS and aRDS adaptation conditions. (**a**) Average vergence and version eye movements for near and far stimuli presented during the cRDS and aRDS adaptation periods. (**b**) Same as (**a**) but for test periods. The dashed lines in (**a**) indicate stimulus onset and offset; coloured shaded regions indicate ±1 s.e.m. and grey shaded bars indicate significant differences between conditions.

### ERP analysis of adaptation stimuli

The primary SSVEP analysis revealed a difference in stereoscopic event-related neural responsiveness following adaptation to aRDS but not cRDS, relative to baseline, consistent with the prediction that adaptation to binocular mismatches increases neural excitability. If the c/aRDS adaptation stimuli have different functional consequences, this indicates that these stimuli evoked different patterns of activity during adaptation. While the central aim of the experiment was to test the consequences of adaptation to c/aRDS, the ERPs evoked by these stimuli during the adaptation period may inform the mechanism by which different effects of adaptation were produced. Thus, in an exploratory analysis, we computed the difference between c/aRDS ERPs (averaged across all presentations, across all sensors). We found that the sensors that showed the greatest difference were regionally similar to those used in the SSVEP analysis (**Fig 5a**). Given the topographic similarity of differences in activity evoked by test and adaptor stimuli between conditions, we then compared the pooled activity of adaptor ERPs from the same (occipital & parietal) sensors used in the previous analysis. We found that the amplitude of the P1 component was significantly smaller in response to aRDS compared to cRDS, but the amplitude of the N1 component was significantly larger (peak t_30_=3.54, P=.001; **Fig 5b**).

**Figure 5.**
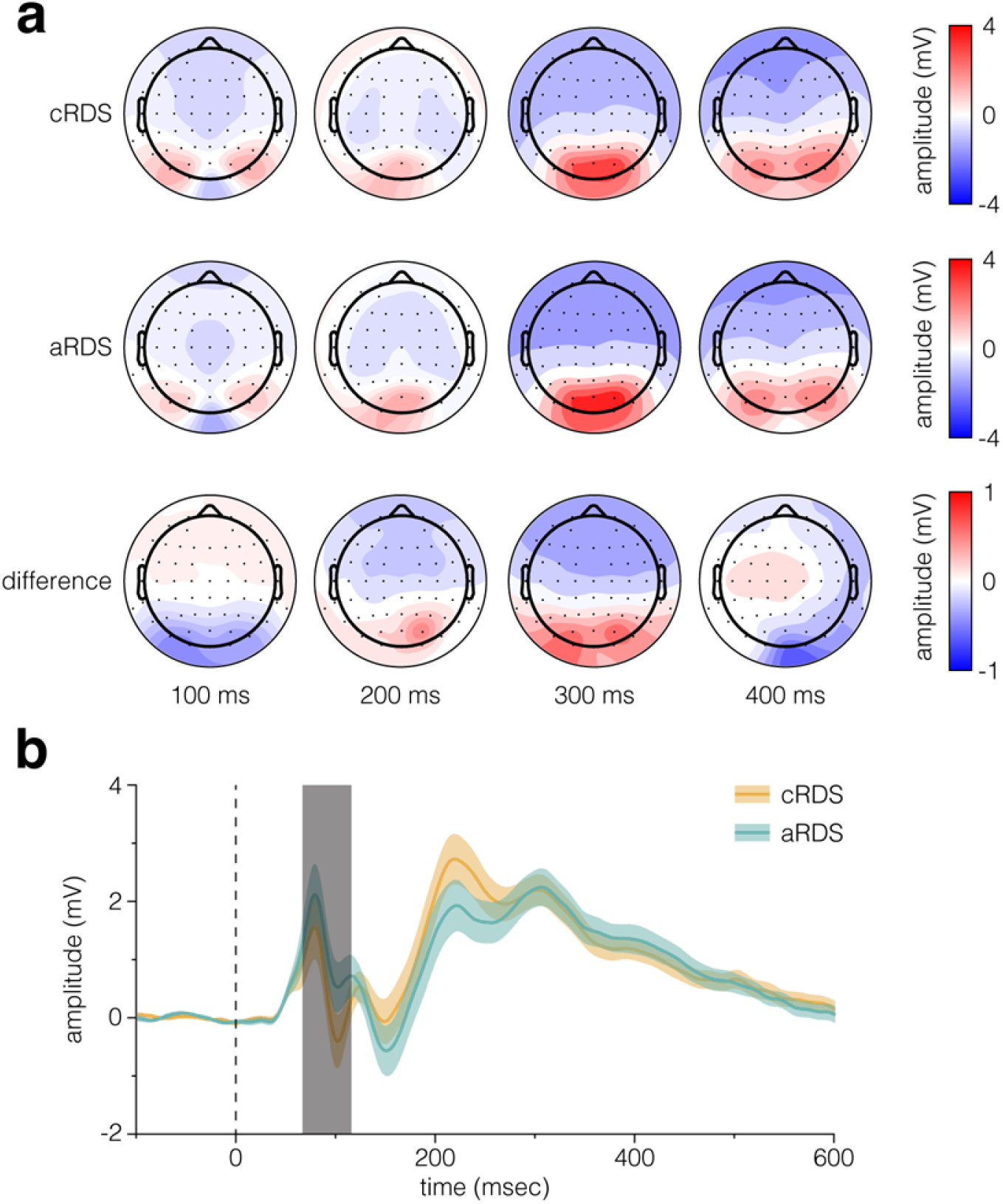
Event-related potentials (ERPs) in response to the adaptation stimuli. **(a)** Topographic maps showing average neural activity in response to the (c/aRDS) adaptation stimuli, and the difference between them, at 100, 200, 300, and 400 ms after stimulus onset. (**b**) Neural activity in response to the (c/aRDS) adaptation stimuli, averaged over the occipital and parietal sensors, as a function of time. The dashed line indicates stimulus onset; coloured shaded regions indicate ±1 s.e.m. and grey shaded bars indicate significant differences between conditions.

**Figure 5.**
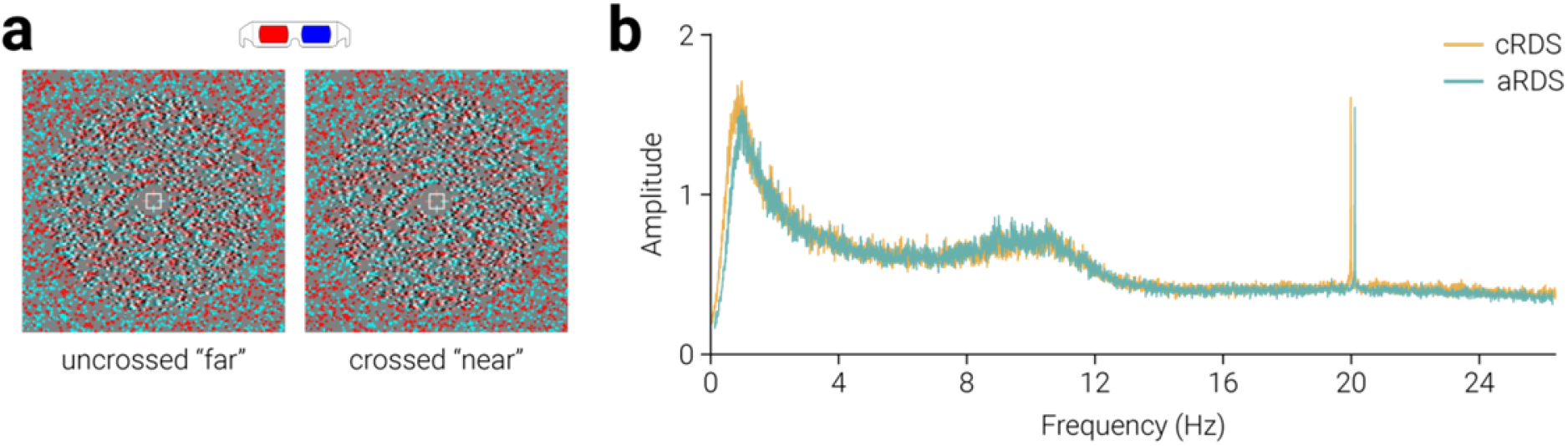
Stimuli and results from the control experiment. **(a)** Example of the aRDS test stimuli used in the control experiment. (b)The SSVEP SNR response spectra (averaged across parietal and occipital sensors) to the aRDS test stimulus following adaptation to c/aRDS, for all participants (*n*=22). The aRDS spectrum is horizontally offset to facilitate comparison with the cRDS spectra.

### Anticorrelated test stimulus

We used a cRDS test stimulus, instead of an aRDS, to measure changes in neural excitability following adaptation to anticorrelated images as cRDS evoke a larger response than aRDS (Braddick et al., 1980; Petrig et al., 1981), providing a better signal-to-noise ratio and thus a more sensitive measure of neural change. However, a possible concern is that the reduction in SSVEP amplitude we observed in response to the change in the depth of a cRDS following adaptation to anticorrelation was influenced by the difference between adaptation and test stimuli. That is, the cRDS test stimulus evoked a reduced neural response following adaptation to aRDS compared to cRDS because the difference between adaptation and test stimulus was more pronounced in the aRDS condition. To test this possibility we repeated the experiment, on a new cohort of participants, using an aRDS test stimulus (**Fig. 6a**).

We included data from the same sensors used in the SSVEP analysis of the main experiment (i.e., Oz, O1, O2, Poz, PO3, PO4, PO7, PO8, Pz, P1, P2, P3, P4, P5, P6, P7, & P8). The SSVEP SNR response spectra for adaptation conditions are shown in **Figure 5b**. While there is a clear peak at the dot refresh frequency (20 Hz), there is little discernible peak at the depth alternation frequency (4 Hz); contrast this with the spectra from the main experiment where a peaks can be clearly resolved at 4 Hz and its harmonics (**Fig. 2c**). Indeed, a comparison of the SSVEP SNR at 4 Hz between the baseline condition here and in the main experiment show that the amplitude was significantly lower here, in response to an aRDS test stimulus (independent t-test: t_43_=5.19, P=5.4e^−6^). We found no significant difference in the 4 Hz SSVEP SNR between aRDS and cRDS adaptation conditions (t_20_=1.21, P=.238). However, if there were differential effects of adaptation to cRDS and aRDS on the potential evoked by the aRDS test stimulus, given the low amplitude of the signal, it is unlikely that this difference could be detected.

## DISCUSSION

Electrophysiological recordings from macaque visual cortex (Cumming & Parker, 1997; Janssen et al., 2003; Tsao et al., 2003) and psychophysical work with humans (Katyal et al., 2018; Kingdom et al., 2018) has revealed the existence of cortical mechanisms tuned to mismatched features between the left and right eyes. While the evidence supporting the existence of neurons tuned to mismatches is extensive, our understanding of their role in binocular vision remains limited. Here we provide evidence that neural activity in the visual cortex may facilitate binocular vision through inhibition. In particular, we show that adaptation to mismatched binocular stimuli, i.e., anticorrelated random-dot stereograms (aRDS), produces increased excitability in the visual cortex in response to changes in depth.

Prolonged/repeated exposure typically produces a reduction in the responsiveness of stimulated neurons, i.e., adaptation. Thus, one might expect adaptation to aRDS to reduce the net responsiveness of the visual cortex in response to a change in depth. Alternatively, one might anticipate that adaptation to aRDS to have no effect on the response to a correlated RDS (cRDS), due to the perceptual dissimilarity of these stimuli, i.e., aRDS do not produce a percept of depth. However, in contrast to these intuitive hypotheses, we found that adaptation to aRDS yields an increase in excitability. While these results may seem surprising, they are consistent with the notion that neurons tuned to binocular mismatches can facilitate stereopsis by suppressing unlikely perceptual interpretations (Goncalves & Welchman, 2017).

The results of the SSVEP analysis of activity evoked by cRDS stimuli in the test period suggested that prolonged viewing of cRDS and aRDS resulted in adaptation of different neural ensembles, or similar neural ensembles adapted to different extents. Analysis of the ERPs evoked by the adaptor stimuli confirmed this, revealing different patterns of activity corresponding to cRDS and aRDS stimuli. Specifically, we found differences between the P1 and N1 components. P1 and N1 components are thought to represent activity relating to early sensory processes; thus, the differences in amplitude of these components in response to aRDS and cRDS may reflect the differences in the engagement of excitatory vs. inhibitory mechanisms. The amplitude of N1, but not P1, has been shown to increase with attentional allocation (Haider, Spong, & Lindsley, 1964; Hillyard & Anllo-Vento, 1998; Polich, 1986; Van Voorhis & Hillyard, 1977). As the amplitude of the N1, but not P1, component was higher for aRDS, this may signal that observers allocated more attention to the aRDS than the cRDS, e.g., because of its perceptual peculiarity. However, the results from the concurrent (attentionally demanding) task suggest otherwise: no differences in accuracy or response time were observed between conditions.

We did not find a reduction in SSVEP amplitude following prolonged exposure to cRDS, which one may have expected based on adaptation. One possible explanation for this is that the correlation between left- and right-eye images in the cRDS is more similar to that observed in the natural environment, compared to the aRDS which has an artificially low correlation. Thus, the tolerance for cRDS may be higher than that for aRDS, and effects of adaptation may be subtler. However, we found a reduction in the amplitude of the ERP over the course of the adaptation period for both aRDS and cRDS, indicating that adaptation had a measurable effect on neural activity for both types of stimuli.

A possible concern is that the increase in SSVEP amplitude following adaptation to aRDS, compared to cRDS, was due to the aRDS stimulus being more perceptually dissimilar to the test stimulus than the cRDS. In particular, the increased response may be due to less expectation of the test stimulus in the aRDS condition. The results from the control experiment, in which we used an aRDS test stimulus, provided inconclusive evidence for this possibility, as the amplitude of the SSVEP SNR was too low to produce reliable estimates. This is consistent with previous work showing cRDS evoke a larger response than aRDS (Braddick et al., 1980; Petrig et al., 1981). However, in the baseline condition of the main experiment, the test stimulus was preceded by a period in which a grey background was presented; this is arguably more dissimilar from the test stimulus than either c/aRDS adaptor. Thus, if SSVEP amplitude reflected the similarity between the test stimuli and preceding images, we would expect SSVEP amplitude to be highest in the baseline condition where the dissimilarity was highest. However, we found no evidence for this: baseline amplitude was significantly lower than that following adaptation to aRDS.

While EEG has relatively high temporal resolution, the spatial resolution of the technique is limited. Thus, a limitation of the current study is that we cannot make precise statements about the likely neural locus of adaptation to binocular anticorrelation in the visual cortex. fMRI is known to have much better spatial resolution compared to EEG, however, excitatory and inhibitory activity cannot be differentiated from BOLD signal, restricting the diagnostic utility of this technique in establishing the role of ‘what not’ mechanisms.

The current results have implications beyond stereopsis. There is theoretical and empirical evidence supporting the existence of neurons tuned to mismatches from studies of stereopsis (DeAngelis, Ohzawa, & Freeman, 1991; Prince, Cumming, & Parker, 2002; Tsao et al., 2003), binocular rivalry (Katyal et al., 2018; Kingdom et al., 2018; Said & Heeger, 2013), and integration of cues within (Kim, Angelaki, & Deangelis, 2015; Nadler et al., 2013; Rideaux & Welchman, 2018) and between sensory modalities (Gu, Angelaki, & DeAngelis, 2008; Kim, Pitkow, Angelaki, & DeAngelis, 2016; Morgan, DeAngelis, & Angelaki, 2008). Here we provide evidence suggesting a role for mechanisms tuned to mismatches that may facilitate inference by driving suppression.

## Competing Interests

The authors declare that they have no competing interests.

## Acknowledgements

We thank N. Goncalves for detailed discussions and comments on the paper. This work was supported by the Leverhulme Trust (ECF-2017-573) and the Wellcome Trust (095183/Z/10/Z).

